# RIPK1 is essential for Herpes Simplex Virus-triggered ZBP1-dependent necroptosis in human cells

**DOI:** 10.1101/2024.09.17.613393

**Authors:** Oluwamuyiwa T. Amusan, Shuqi Wang, Chaoran Yin, Heather S. Koehler, Yixun Li, Tencho Tenev, Rebecca Wilson, Benjamin Bellenie, Ting Zhang, Jian Wang, Chang Liu, Kim Seong, Seyedeh L. Poorbaghi, Joseph Yates, Yuchen Shen, Jason W. Upton, Pascal Meier, Siddharth Balachandran, Hongyan Guo

## Abstract

Necroptosis initiated by the host sensor Z-NA Binding Protein-1 (ZBP1) is essential for host defense against a growing number of viruses, including Herpes Simplex Virus-1 (HSV-1). Studies with HSV-1 and other necroptogenic stimuli in murine settings have suggested that ZBP1 triggers necroptosis by directly complexing with the kinase RIPK3. Whether this is also the case in human cells, or whether additional co-factors are needed for ZBP1-mediated necroptosis, is unclear. Here, we show that ZBP1-induced necroptosis in human cells requires RIPK1. We have found that RIPK1 is essential for forming a stable and functional ZBP1-RIPK3 complex in human cells, but is dispensable for the formation of the equivalent murine complex. The RIP Homology Interaction Motif (RHIM) in RIPK3 is responsible for this difference between the two species, because replacing the RHIM in human RIPK3 with the RHIM from murine RIPK3 is sufficient to overcome the requirement for RIPK1 in human cells. These observations describe a critical mechanistic difference between mice and humans in how ZBP1 engages in necroptosis, with important implications for treating human diseases.

## Introduction

Necroptosis contributes to both antiviral host defense and inflammatory tissue damage [1-3]. Necroptosis is triggered when receptor-interacting protein (RIP) kinase (RIPK)3 phosphorylates and activates the pseudo kinase mixed-lineage kinase domain-like (MLKL) [4-7], resulting in membrane permeabilization and necrotic death [8-11]. In humans and mice, RIPK3 may be activated by one of three different adaptors that contain at least one RIP homotypic interaction motif (RHIM): (i) RIPK1 downstream of TNF family death receptors [4, 5]; (ii) TIR domain-containing adapter-inducing interferon (TRIF) downstream of Toll-like receptor (TLR)3 or TLR4 [12, 13]; or (iii) Z-form nucleic acid (Z-NA)-binding protein 1 (ZBP1, also known as DAI or DLM-1) [14, 15].

ZBP1 was initially described as an interferon (IFN) induced protein [16], and proposed to sense cytosolic dsDNA and activate a type I IFN response [17]. Subsequent studies, however, showed that ZBP1 was largely dispensable for inducing Type I IFNs downstream of dsDNA [18]. Later, ZBP1 was found to contain a functional RHIM and shown to bind RIPK3, indicative of a potential upstream role in RIPK3-driven necroptosis [19]. ZBP1 has since been demonstrated to mediate necroptosis activation during viral infections. Although first observed with murine cytomegalovirus (MCMV) [15], ZBP1-driven cell death has been reported in cells infected with influenza A and B viruses (IAV and IBV) [20, 21], orthopoxvirus (vaccinia virus, VACV) [22], herpes simplex virus-1 (HSV-1) [23], and SARS-CoV-2 [24-26]. Recently, ZBP1 has been shown to recognize Z-RNA as the ligand to trigger RIPK3-mediated necroptosis during virus infection [27, 28]. ZBP1-dependent necroptosis not only curtails viral replication and limits dissemination but also contributes to severe inflammatory tissue damage [29, 30].

Virus-triggered ZBP1-induced RIPK3 activation in mouse cells does not require RIPK1. For example, during MCMV infection of mouse cells, ZBP1 directly recruits RIPK3 and induces necroptosis without the need for RIPK1 [15, 31, 32]. HSV-1 and IAV infections also induce ZBP1-driven necroptosis in mouse cells that is independent of RIPK1 [20, 21, 23]. Thus, it is commonly assumed that once ZBP1 binds Z-form nucleic acid ligands during viral infection, it directly associates with and activates RIPK3 [29]. Whether ZBP1 directly activates RIPK3 in human cells is unknown.

Using a necroptogenic HSV-1 mutant virus [23], we show that ZBP1-driven necroptosis in human cells requires RIPK1. ZBP1 binds to RIPK1 to efficiently recruit RIPK3 and executes necroptosis in infected human cells. Mechanistically, we determine that RIPK1 stabilizes the human ZBP1-RIPK3 complex and promotes its polymerization into functional detergent-insoluble amyloid structures. In contrast, RIPK1 is not essential for the ZBP1-RIPK3 association or the initiation of virus-induced necroptosis in mouse cells. The murine ZBP1-RIPK3 complex forms amyloid fibrils and activates necroptosis independent of RIPK1. Replacing the RHIM in human RIPK3 with its murine counterpart is sufficient to bypass the need for RIPK1 in human cells. Overall, our results unmask a species-specific requirement for RIPK1 in promoting ZBP1-RIPK3 signaling in human cells. These results now position RIPK1 inhibitors, already in advanced clinical trials for a range of human diseases, as potential therapeutics for ZBP1-initiated inflammatory conditions as well.

## Results

### HSV-1(ICP6mut) infection triggers endogenous ZBP1-driven necroptosis in human cells

HSV-1 encodes an RHIM-containing protein, ICP6. When this RHIM is rendered non-functional by mutation, HSV-1 triggers ZBP1-dependent necroptosis in both murine and human cells [23]. The HSV-1(ICP6mut) virus is thus an excellent tool with which to dissect ZBP1-initiated necroptosis in both species. To explore ZBP1-induced signaling in human cells, we utilized a cell line (HT-29) that expresses RIPK3 and MLKL, and in which endogenous ZBP1 can be induced by IFN-β pre-treatment. We then used CRIPSR-Cas9 technology to ablate ZBP1 in these cells. After confirming the knockout (KO) by sequencing and immunoblotting (Fig.1A and B), we primed these cells with IFN-β for 16 hrs, and infected them with HSV-1(ICP6mut). HSV-1(ICP6mut) efficiently triggered cell death in the ZBP1-expressing parental cells (Fig. 1C), associated with the phosphorylation of RIPK1, RIPK3, and MLKL (Fig.1D), but not in the ZBP1 KO cells. We also found that HSV-1(ICP6mut) induced ZBP1-dependent cell death in primary human foreskin fibroblasts (HFFs) (Fig. 1E and F). Altogether, these findings support the idea that HSV-1(ICP6mut) infection triggers ZBP1-driven necroptosis in human cells.

**Figure 1.**
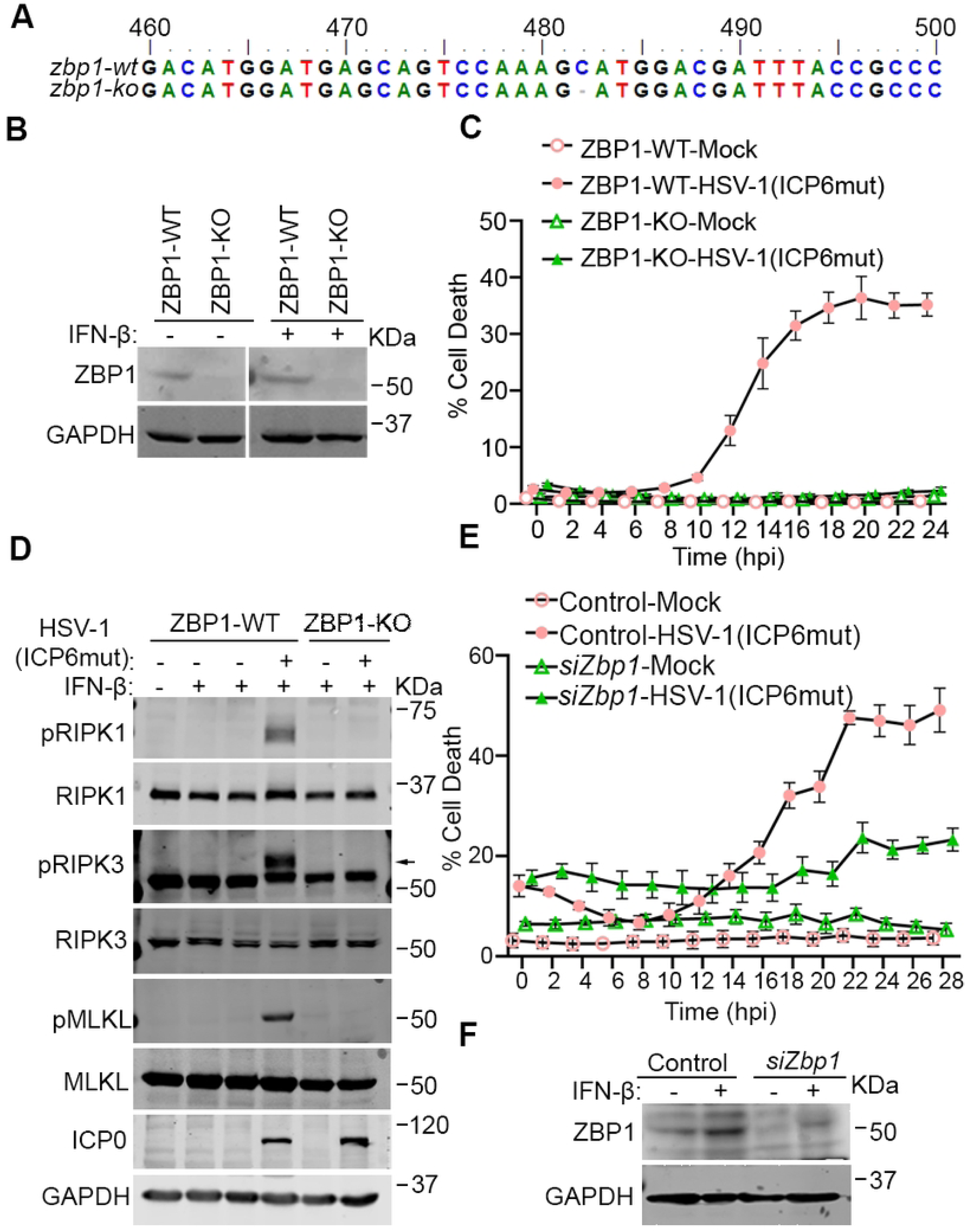
HSV-1(ICP6mut) infection triggers endogenous ZBP1-driven necroptosis in human cells. (A) Genomic DNA sequences of ZBP1 locus in ZBP1 KO HT29 cells. ‘-’ indicates none of the nucleotides. (B) Immunoblotting analysis of endogenous expression of Wild-type (WT) or ZBP1 KO HT29 cells with or without IFN-β priming. (C) Cell death kinetics of WT or ZBP1 KO HT29 cells after IFN-β priming, followed by HSV-1(ICP6mut) infection. (D) Immunoblotting of WT or ZBP1 KO HT29 cells with or without IFN-β priming, followed by HSV-1(ICP6mut) infection. Cells were collected after 16hrs infection and subjected to immunoblotting analysis with pRIPK1, RIPK1, pRIPK3, RIPK3, p-MLKL, MLKL, ICP0, and GAPDH antibodies. (E) Cell death kinetics of primary human foreskin fibroblast (HFF) cells transfected with control or ZBP1 siRNAs using electroporation and primed with IFN-β (50ng/ml) for 24hrs, followed by HSV-1(ICP6mut) infection. The knockdown efficiency in HFFs was confirmed by immunoblotting assay in (F). Results are representative of at least two independent experiments. Error bars represent mean ± SD.

### RIPK1 is essential for HSV-1(ICP6mut)-triggered endogenous ZBP1-driven necroptosis in human cells

Expectedly, treatment of HT-29 cells with the RIPK3 kinase inhibitor GSK872 or the MLKL inhibitor NSA, but not the pan-caspase inhibitor zVAD, was sufficient to prevent ZBP1-mediated cell death triggered by HSV-1(ICP6mut) (Fig. 2B), consistent with our previous findings [23]. Surprisingly, inhibiting RIPK1 kinase activity with Nec-1 also efficiently blocked HSV-1(ICP6mut)-triggered MLKL phosphorylation and cell death. (Fig. 2B and C). To investigate the unanticipated dependency of RIPK1 for ZBP1-mediated necroptosis in human cells, we utilized a proteolysis-targeting chimera (PROTAC) that harnesses the ubiquitin-proteasome system to transiently and efficiently degrade RIPK1 (Fig.2A). The RIPK1 PROTAC R1-ICR-3 efficiently degraded RIPK1 protein in all human (and murine) cell types tested [33]. Consistent with observations using the RIPK1 kinase inhibitor, R1-ICR-3 blocked HSV-1(ICP6mut) triggered-cell death (Fig.2B). RIPK1, RIPK3, and MLKL phosphorylation levels were also reduced in HSV-1(ICP6mut)-infected cells treated with R1-ICR-3 (Fig.2C).

**Figure 2.**
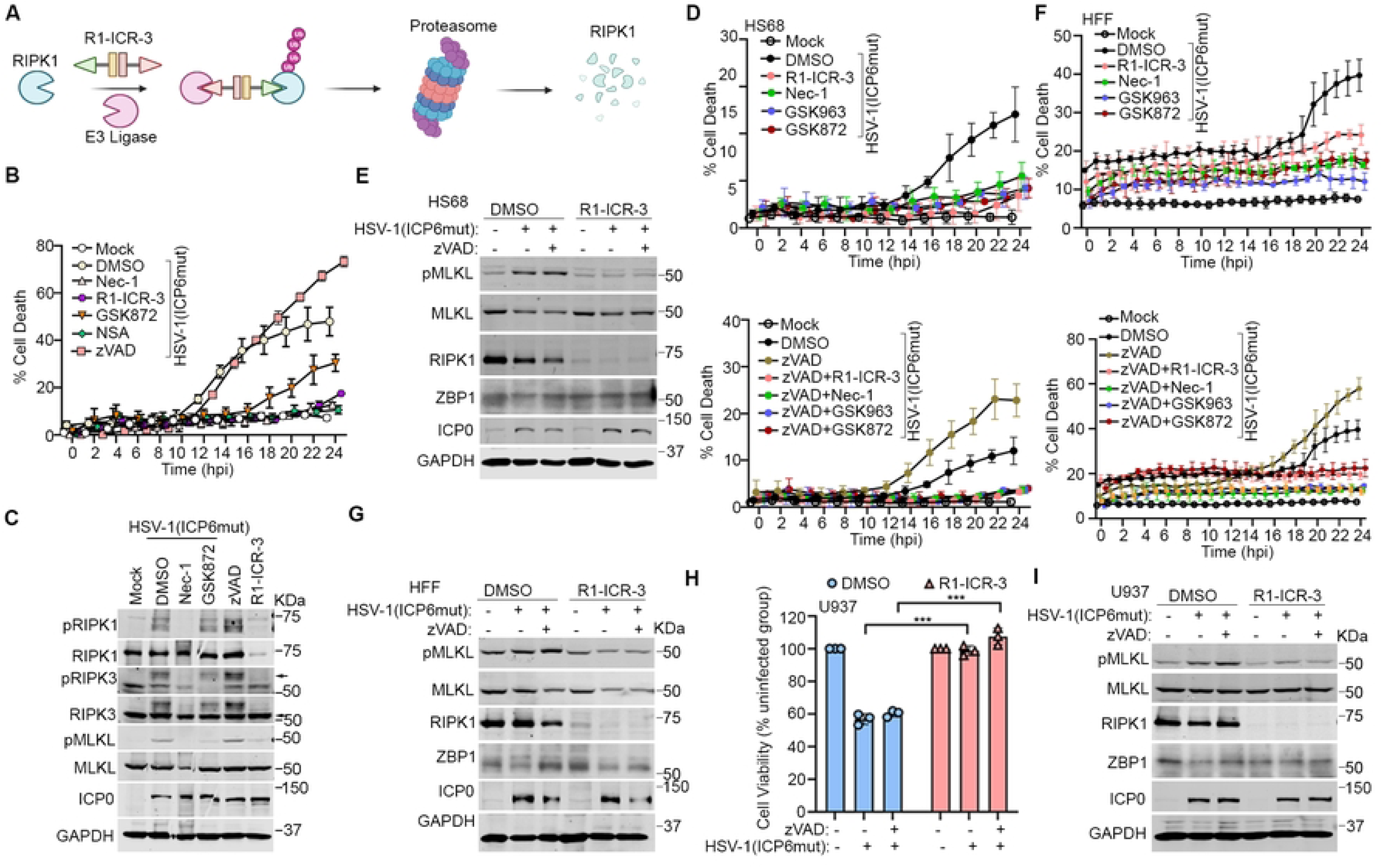
RIPK1 is essential for HSV-1(ICP6mut)-triggered endogenous ZBP1-driven necroptosis in human cells. (A) Schematic illustration of the PROTAC-induced RIPK1 degradation system. (B) Cell death kinetics of HT29 cells pretreated with IFN-β (50 ng/mL) for 24 hours, followed by HSV-1(ICP6mut) infection, with or without Nec-1 (30 µM), GSK872 (5 µM), NSA (2 µM), R1-ICR-3 (100 nM), or zVAD (25 µM). (C) Immunoblotting of HT29 cells primed with IFN-β (50 ng/mL), followed by HSV-1(ICP6mut) infection, with or without Nec-1 (30 µM), GSK872 (5 µM), NSA (2µM), zVAD (25 µM), or R1-ICR-3 (100 nM). Cells were subjected to immunoblotting with pRIPK1, RIPK1, pRIPK3, RIPK3, pMLKL, MLKL, ICP0, and GAPDH antibodies at 16 hours post-infection (hpi). (D) Cell death kinetics of HS68 cells pretreated with IFN-β (50 ng/mL) for 24 hours, followed by HSV-1(ICP6mut) infection, with or without zVAD (25 µM), in combination with Nec-1 (30 µM), GSK963 (5 µM), GSK872 (5 µM), or R1-ICR-3 (100 nM). (E) Immunoblotting of HS68 cells pretreated with IFN-β (50ng/ml) for 24 hrs, followed by HSV-1(ICP6mut) infection, with or without zVAD (25 µM), in combination with R1-ICR-3 (100 nM). Cells were collected at 16 hours post-infection and probed with pMLKL, MLKL, RIPK1, ZBP1, ICP0, and GAPDH antibodies. (F) Cell death kinetics of primary human foreskin fibroblast (HFF) cells pretreated with IFN-β (50 ng/mL) for 24 hours, followed by HSV-1(ICP6mut) infection, with or without zVAD (25 µM), in combination with Nec-1 (30 µM), GSK963 (5 µM), GSK872 (5 µM), or R1-ICR-3 (100 nM). (G) Immunoblotting of HFF cells pretreated with IFN-β (50ng/ml) for 24 hrs, followed by HSV-1(ICP6mut) infection, with or without zVAD (25 µM), in combination with R1-ICR-3 (100 nM). Cells were collected at 16 hpi and probed with pMLKL, MLKL, RIPK1, ZBP1, ICP0, and GAPDH antibodies. (H) Cell viability of U937 cells, followed by HSV-1(ICP6mut) infection, with or without zVAD (25 µM), in combination with R1-ICR-3 (100 nM) at 18 hpi. (I) Immunoblotting of U937 cells, followed by HSV-1(ICP6mut) infection, with or without zVAD (25 µM), in combination with R1-ICR-3 (100 nM). Samples were probed for p-MLKL, MLKL, RIPK1, ZBP1, ICP0, and GAPDH antibodies at 12 hpi. Results are representative of at least two independent experiments. Error bars represent mean ± SD.

We then sought to confirm whether RIPK1 is essential for HSV-1(ICP6mut)-triggered ZBP1-mediated necroptosis in other human cell types. In IFN-β-primed HS68 cells (Fig.2D) and HFFs (Fig.2F), HSV-1(ICP6mut)-triggered cell death was prevented by the RIPK1 degrader R1-ICR-3, RIPK1 kinase inhibitors including Nec-1 and GSK963, and the RIPK3 kinase inhibitor GSK872, but not by the pan-caspase zVAD. RIPK1 degradation with R1-ICR-3 also reduced MLKL phosphorylation during HSV-1(ICP6mut) infection in both HS68 (Fig.2E) and HFFs (Fig.2G). We observed similar results in human U937 monocytes, where HSV-1(ICP6mut)-induced cell death was effectively prevented by R1-ICR-3 (Fig.2H), accompanied by a reduction in MLKL phosphorylation (Fig.2I). Taken together, these findings demonstrate an unexpectedly essential role for RIPK1 in ZBP1-mediated necroptosis signaling in human cells.

Importantly, RIPK1 degradation did not rescue murine cells from HSV-1-induced death (Fig.S1A-F). Although RIPK1 kinase inhibition delayed HSV-1(ICP6mut)-induced necroptosis, it still resulted in comparable cell death levels at later time points (Fig.S1G). RIPK1 thus does not play an obligatory role in HSV-1(ICP6mut)-triggered necroptosis in mouse cells.

### RIPK1 is recruited to the ZBP1-RIPK3 complex in human cells during HSV-1(ICP6mut) infection

To explore the mechanistic basis for why RIPK1 was needed for ZBP1-induced cell death in human cells, we appended an N-terminal FLAG tag to ZBP1 and stably expressed this construct in HT-29 cells (hereafter HT29-hZBP1) [23, 27]. HSV-1(ICP6mut)-infected HT29-hZBP1 cells failed to undergo necroptosis when treated with the RIPK1 degrader R1-ICR-3, RIPK1 kinase inhibitors Nec-1 and GSK963, or the RIPK3 kinase inhibitor GSK872 (Fig.3A), similar to what we observed with endogenous ZBP1 (Fig.2B). Activation markers of ZBP1-mediated necroptosis, including RIPK3 phosphorylation (pRIPK3) and MLKL phosphorylation (pMLKL), were attenuated in R1-ICR-3-treated HT29-hZBP1 cells during HSV-1(ICP6mut) infection (Fig.3B). These findings demonstrate that FLAG-tagged ZBP1 functions similarly to endogenous ZBP1 and that RIPK1 is essential in ZBP1-mediated necroptosis.

**Figure 3.**
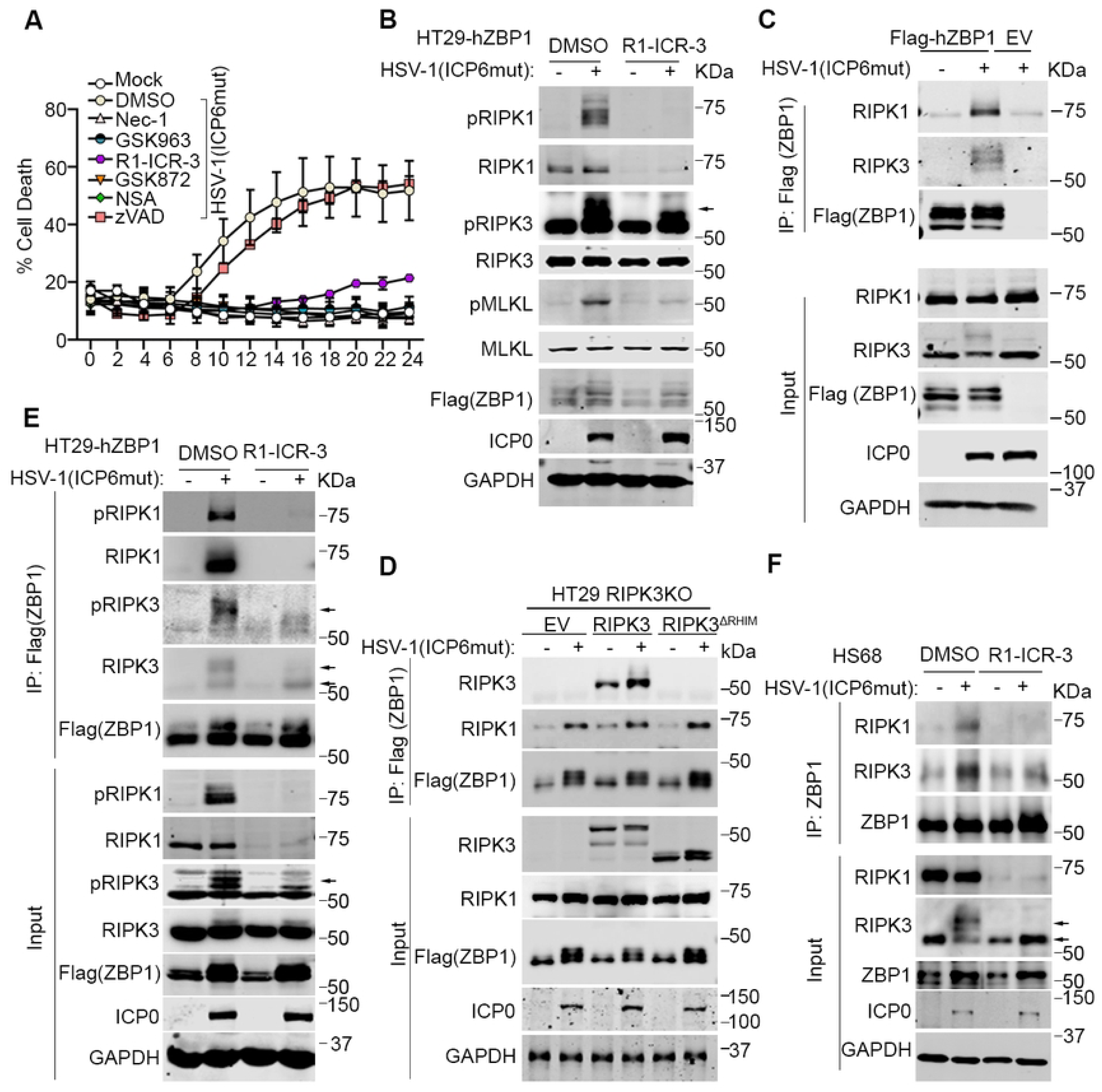
RIPK1 binds to ZBP1 during HSV-1(ICP6mut) infection in human cells. (A) Cell death kinetics of HT29 cells reconstituted Flag-hZBP1 (hereafter HT29-hZBP1), followed by HSV-1(ICP6mut) infection, with or without Nec-1 (30 µM), GSK963 (5 µM), GSK872 (5 µM), NSA (2 µM), R1-ICR-3 (100 nM), or zVAD (25 µM). (B) Immunoblotting of HT29-hZBP1 cells, followed by HSV-1(ICP6mut) for 10 hrs with or without R1-ICR-3 (100 nM). Samples were probed with pRIPK1, RIPK1, pRIPK3, RIPK3, pMLKL, MLKL, Flag (ZBP1), ICP0, and GAPDH antibodies. (C) HT29-hZBP1 cells either mocked or infected with HSV-1(ICP6mut) for 10 hrs were harvested and subjected to immunoprecipitation with Anti-FLAG® M2 magnetic beads (Sigma-Aldrich). Total lysates (Input) and immunoprecipitates (IP) were subjected to SDS-PAGE and evaluated for RIPK1, RIPK3, and Flag (ZBP1). Input was also evaluated for viral protein ICP0 and GAPDH. (D) HT29 RIPK3 knockout (KO) cells reconstituted empty vector (EV), wild-type (WT) RIPK3, or RHIM-deleted (ΔRHIM) RIPK3 together with human ZBP1, were either mock or infected with HSV-1(ICP6mut) for 10 hrs. Co-immunoprecipitation was performed in these cells as described in C. (E) HT29-hZBP1 cells either mocked or infected with HSV-1(ICP6mut) for 10 hrs with or without R1-ICR-3 (100 nM), were harvested and subjected to immunoprecipitation as described in C. (F) HS68 cells pretreated with IFN-β (50ng/ml) for 24 hrs, followed by HSV-1(ICP6mut) infection in the presence of zVAD (25 µM), with or without R1-ICR-3 (100 nM) for 16 hrs, were harvested and subjected to immunoprecipitation as described in C. Results are representative of at least two independent experiments. Error bars represent mean ± SD.

To determine whether RIPK1 binds to ZBP1 during viral infection, we used the FLAG antibody to immunoprecipitate FLAG-ZBP1 from infected HT29-hZBP1 cells. Mock-infected HT29-hZBP1 and viral-infected HT29-EV cells served as controls. RIPK1, along with RIPK3, was detected in ZBP1 complexes isolated from infected HT29-hZBP1 cells but not from the control groups (Fig.3C). In contrast, only trace amounts of RIPK1 co-immunoprecipitated with ZBP1 in mouse cells (Fig.S1H), consistent with previous reports [34]. As RIPK1 can be recruited to ZBP1 via RIPK3 during virus-induced apoptosis [35], we next investigated whether the recruitment of RIPK1 to ZBP1 depended on RIPK3. We used CRISPR-based approaches to knock out (KO) RIPK3 in HT29 cells, reconstituted the cells with empty vector (EV), wild-type (WT) RIPK3, or RIPK3 (ΔRHIM) [36], and infected them with HSV-1(ICP6mut). As shown in Fig.3D, RIPK1 was immunoprecipitated from extracts of HT29 (RIPK3 KO) cells reconstituted with EV as well as RIPK3 (ΔRHIM) cells after viral infection, despite the absence of RIPK3 or the inability to participate in RHIM-dependent interactions, demonstrating that recruitment of RIPK1 to ZBP1 is independent of RIPK3. Instead, we found that RIPK1 was needed for RIPK3 recruitment to hZBP1. RIPK1 degradation with R1-ICR-3 treatment prevented RIPK3, especially activated RIPK3 (pRIPK3), from binding to ZBP1 during HSV-1(ICP6mut) infections (Fig.3E). Similarly, R1-ICR-3 also reduced RIPK3 recruitment by endogenous ZBP1 triggered by HSV-1(ICP6mut) infection in HS68 cells (Fig.3F). Taken together, these findings indicate that hZBP1 recruits RIPK1 before RIPK3, and RIPK1 functions as an essential adaptor for the formation of the ZBP1-RIPK3 complex in HSV-1(ICP6mut)-infected human cells.

To further confirm the role of RIPK1 in ZBP1-driven necroptosis, we used the CRISPR method to knock out RIPK1 in HT29 cells and then reconstituted with FLAG-tagged human ZBP1 (Flag-hZBP1). RIPK1 knockout prevented HSV-1(ICP6mut)-triggered cell death (Fig.4A), consistent with the effects observed with RIPK1 degrader or kinase inhibitor treatments (Fig.3A). Additionally, RIPK3 and MLKL phosphorylation were also inhibited in these RIPK1 KO cells (Fig.4B). To investigate whether the RIPK1 dependency is due to species-specific differences, RIPK1 KO cells were reconstituted with either human or mouse RIPK1, generously provided by Dr. Pascal Meier. As expected, reconstitution with either human or mouse RIPK1 in RIPK1 KO cells restored RIPK1-dependent apoptosis induced by TNF plus Smac mimic (BV-6) treatment, compared to parental RIPK1 KO cells. However, TNF plus CHX could still induce RIPK1-independent apoptosis in all these cells (Fig.4C-E). Interestingly, either murine or human RIPK1 was able to promote the ZBP1-RIPK3 association in human cells, as HSV-1(ICP6mut) infection triggered cell death in RIPK1 KO cells reconstituted with either human or mouse RIPK1 (Fig. 4F). This was associated with the activation of necroptosis markers (pRIPK3 and pMLKL) (Fig.4G).

**Figure 4.**
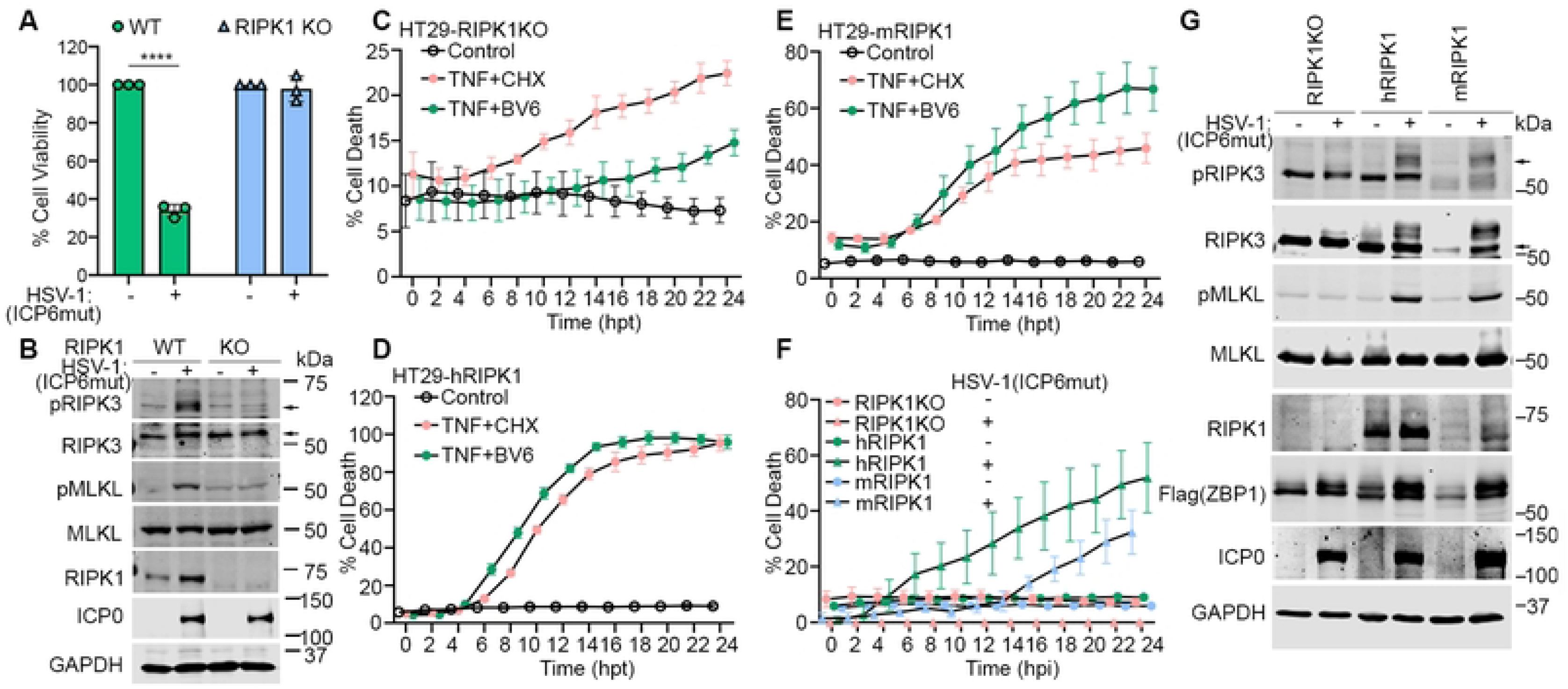
Both human and mouse RIPK1 mediate ZBP1-driven necroptosis during HSV-1(ICP6mut) infection in human cells. (A) Cell viability of HT29-hZBP1 or HT29-hZBP1(RIPK1 KO) cells infected with HSV-1(ICP6mut). Viability was determined at 18 hpi by CellTiter-Glo assay. (B) HT29-hZBP1 or HT29-hZBP1 (RIPK1 KO) cells infected with HSV-1(ICP6mut) were harvested at 10 hpi and subjected to immunoblotting with p-RIPK3, RIPK3, p-MLKL, MLKL, ICP0, and GAPDH antibodies. (C-E) Cell death kinetics of HT29-hZBP1 (RIPK1 KO) (C), HT29-hZBP1(hRIPK1) (D), and HT29-hZBP1(mRIPK1) (E) cells pretreated with Cumate (30µg/ml) for 3 hrs, followed by treatment of TNF+CHX or TNF+BV6. (F) Cell death kinetics of HT29-hZBP1 (RIPK1 KO), HT29-hZBP1(hRIPK1), and HT29-hZBP1(mRIPK1) cells pretreated with Cumate (30µg/ml) for 16 hrs, followed by HSV-1(ICP6mut) infection. (G) Immunoblotting of HT29-hZBP1 (RIPK1 KO), HT29-hZBP1 (hRIPK1), and HT29-hZBP1 (mRIPK1) cells pretreated with Cumate (30µg/ml) for 16hrs, followed by HSV-1(ICP6mut) infection. Cells were harvested at 10 hpi and subjected to immunoblotting with p-RIPK3, RIPK3, p-MLKL, MLKL, RIPK1, Flag (ZBP1), ICP0, and GAPDH antibodies. An unpaired t-test was used to test for statistical differences between indicated conditions in (A). *P < 0.1, **P < 0.0001, ***P < 0.001, ****P < 0.0001. Individual data points indicate three technical replicates. Results are representative of at least two independent experiments. Error bars represent mean ± SD.

### The recruitment of RIPK1 depends on Z**α**2 and RHIM A domains of human ZBP1

ZBP1 has two Z-NA binding domains as well as two RHIMs (Fig.5A). To investigate which of these are essential for HSV-1(ICP6mut)-triggered necroptosis in human cells, we constructed a serial of ZBP1 mutations and reconstituted HT29 cells. HT29 cells expressing hZBP1 mutant Zα1 (mutZα1), or mutant RHIM B (mutRHIMB) showed comparable susceptibility to necroptosis as WT ZBP1, while mutant Zα2 (mutZα2), mutant Zα1Zα2 (mutZα1Zα2), and mutant RHIM A (mutRHIMA) demonstrated resistance to HSV-1(ICP6mut)-induced necroptosis (Fig.5B). Next, we tested if Zα2 and RHIM A in ZBP1 were required for RIPK1 recruitment to ZBP1. Indeed, consistent with the cell viability assay, RIPK1 co-immunoprecipitated with ZBP1 and RIPK3 in necroptosis-susceptible cells (HT29 WT, mutZα1, and mutRHIMB) following viral infection but was not in cells with inactivating mutations in Zα2 or RHIM A (mutZα2, mutZα1Zα2, and mutRHIMA) (Fig.5C). These results suggested that ligand-dependent activation of ZBP1 through Zα2 domain was crucial to recruit RIPK1 via the RHIM A domain.

**Figure 5.**
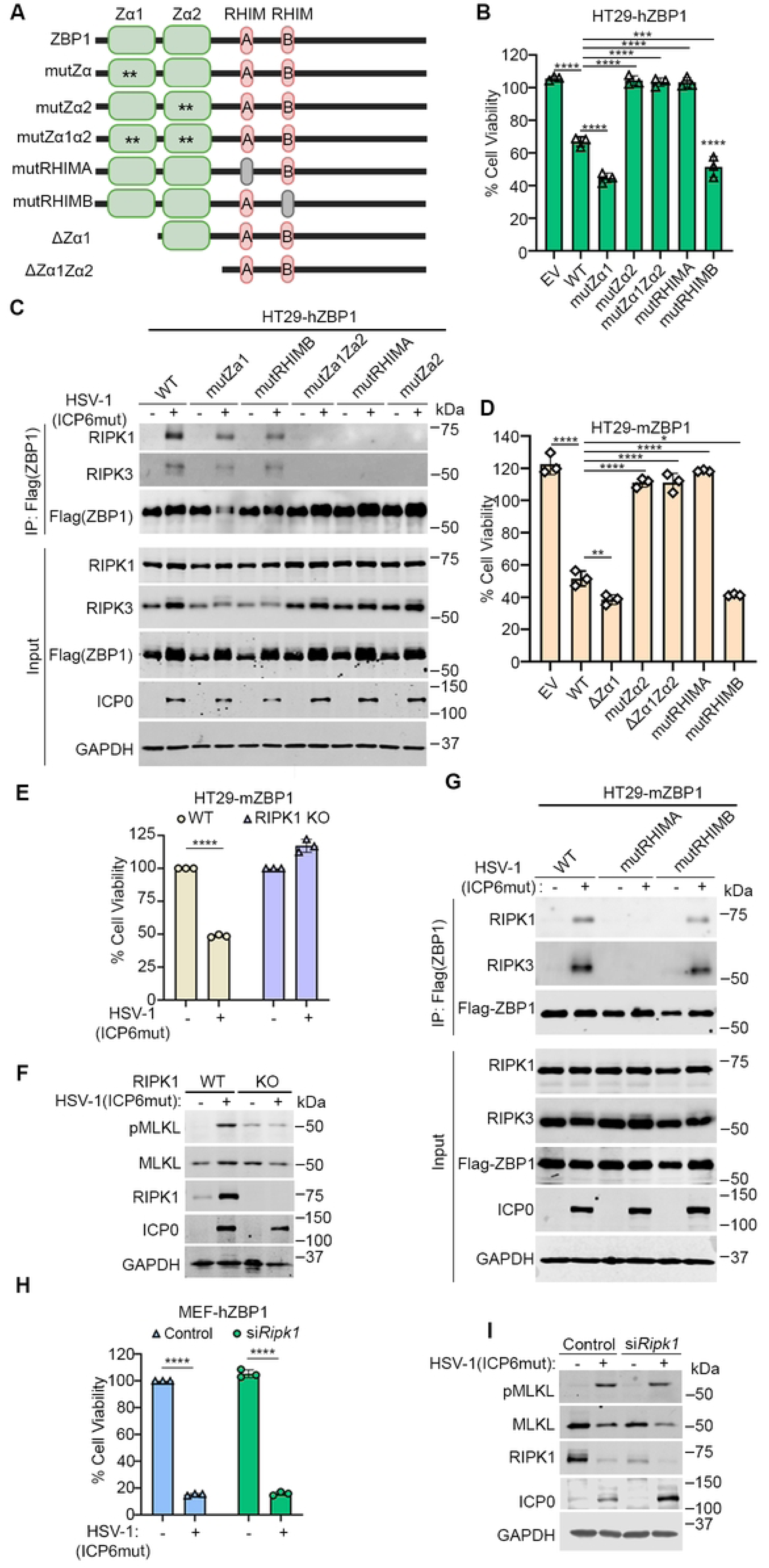
Both Za2 and RHIM A domains of human ZBP1 are essential for RIPK1 binding and mediating necroptosis. (A) Schematic of ZBP1 and its mutants. (B) Cell viability of HT29 cells reconstituted with indicated human ZBP1 constructs infected with HSV-1(ICP6mut). Viability was determined at 18 hpi by CellTiter-Glo assay. (C) HT29 cells reconstituted with indicated human ZBP1 constructs were either mocked or infected with HSV-1(ICP6mut) for 10 hrs. Co-immunoprecipitation was performed in these cells. (D) Cell viability of HT29 cells reconstituted with indicated mouse ZBP1 constructs infected with HSV-1(ICP6mut). Viability was determined at 18 hpi by CellTiter-Glo assay. (E) Cell viability of HT29-mZBP1 or HT29-mZBP1 (RIPK1 KO) cells infected with HSV-1(ICP6mut). Viability was determined at 18 hpi by CellTiter-Glo assay. (F) HT29-mZBP1 or HT29-mZBP1 (RIPK1 KO) cells infected with HSV-1(ICP6mut) were harvested at 10 hpi and subjected to immunoblotting with p-MLKL, MLKL, RIPK1, ICP0, and GAPDH antibodies. (G) HT29 cells reconstituted with indicated mouse ZBP1 constructs were mock-treated or infected with HSV-1(ICP6mut) for 10 hrs. Co-immunoprecipitation was performed in these cells. (H) Cell viability of MEF-hZBP1 transfected with scramble or RIPK1 siRNA for 48 hrs followed by HSV-1(ICP6mut) infection. Viability was determined at 18 hpi by CellTiter-Glo assay. (I) MEF-hZBP1 cells transfected with either scramble siRNA (Control) or RIPK1 siRNA for 48 hrs, followed by HSV-1(ICP6mut) infection. Cells were harvested at 10 hpi and subjected to IB with p-MLKL, MLKL, RIPK1, ICP0, and GAPDH antibodies. One-way ANOVA and Dunnett’s multiple comparisons tests were used to test for statistical differences between indicated conditions and EV groups in (B) and (D). An unpaired t-test was used to test for statistical differences between indicated conditions in (E) and (H). *P < 0.1, **P < 0.0001, ***P < 0.001, ****P < 0.0001. Individual data points indicate three technical replicates. Results are representative of at least two independent experiments. Error bars represent mean ± SD.

Having shown that the human ZBP1-RIPK3 interaction was uniquely dependent on RIPK1, and having ruled out species-specific differences in RIPK1 as the cause for this dependency in human cells, we next asked if species-specific differences in ZBP1 were instead responsible for why only human cells were reliant on RIPK1 for association with RIPK3. To this end, we transduced mouse ZBP1 (mZBP1) into HT29 cells, and found that mZBP1 was able to fully restore susceptibility to HSV-1(ICP6mut)-triggered necroptosis in human cells (Fig.5D). Notably, HT29 transduced with mZBP1ΔZα1, and mutRHIMB showed comparable susceptibility to WT mZBP1 to virus-induced necroptosis, whereas HT29 cells transduced with mZBP1 mutZα2, ΔZα1Zα2, and mutRHIMA showed resistance to virus-induced necroptosis (Fig.5D), in line with our discoveries with hZBP1. We also examined the essential domains of mZBP1 for virus-induced necroptosis in MEFs (Fig.S2A-C). MEFs transduced with mZBP1 WT, ΔZα1, and mutRHIMB but not mutZα2, ΔZα1Zα2, and mutRHIMA were susceptible to virus-induced necroptosis. These results confirm that Zα2 and RHIM A domains are essential for ZBP1-driven necroptosis in mouse cells.

Ablating endogenous hRIPK1 prevented virus-induced necroptosis in HT29-mZBP1 cells (Fig.5E). The necroptosis activation marker (pMLKL) was readily detected in HT29-mZBP1 cells infected by HSV-1(ICP6mut) but not in RIPK1 KO cells (Fig.5F). In agreement with these results, both RIPK1 and RIPK3 co-immunoprecipitated with ZBP1 in HT29-mZBP1 and HT29-mZBP1(mutRHIMB) cells, but not in the death-resistant HT29-mZBP1(mutRHIMA) cells (Fig.5G). In corollary studies, we asked if hZBP1 could substitute for mZBP1 in murine cells. To assess this, we stably expressed hZBP1 in immortalized *Zbp1*^-/-^ MEFs, and found that MEFs transduced with hZBP1 were susceptible to virus-induced necroptosis (Fig.5H). Importantly, RIPK1 was not required for hZBP1 signaling in murine cells (Fig.5I), consistent with the dispensable role of RIPK1 for virus-triggered necroptosis in murine cells (Fig.S1E). In summary, these findings demonstrate that the species-specific requirement for RIPK1 in human cells is not due to differences between human and mouse ZBP1.

### Species-Specific RIPK3 RHIM domain determines the requirement of RIPK1 in virus-triggered ZBP1-dependent necroptosis

Next, we asked if species-specific differences in RIPK3 were why RIPK1 was needed downstream of ZBP1 in human cells. We utilized AlphaFold to model the structures of hZBP1-hRIPK3 and mZBP1-mRIPK3 complexes. Structural comparison of the two ZBP1-RIPK3 complexes revealed several residue variations between the RHIMs of mRIPK3 and hRIPK3 that may result in the different binding affinities to ZBP1 (Fig. 6A and B). Specifically, in the mRIPK3 complex, F442 is fully buried in a hydrophobic pocket and surrounded by the aliphatic side chains of I166, N173, I175, and I184 of mZBP1 (Fig. 6B, left panel). In contrast, Ile452 (I452), the equivalent residue in hRIPK3, was predicted to only form a hydrophobic interaction with Leu401 (L401) of hZBP1, primarily due to its shorter side chain (Fig. 6B, right panel). Furthermore, the three hydrogen bonds mediated by N443, E447, and N452 of mRIPK3 in the mZBP1-mRIPK3 complex (Fig. 6B, left panel) are absent in the hZBP1-hRIPK3 complex, where they are instead substituted with Y453, G457, and D462 in hRIPK3 (Fig. 6B, right panel).

**Figure 6.**
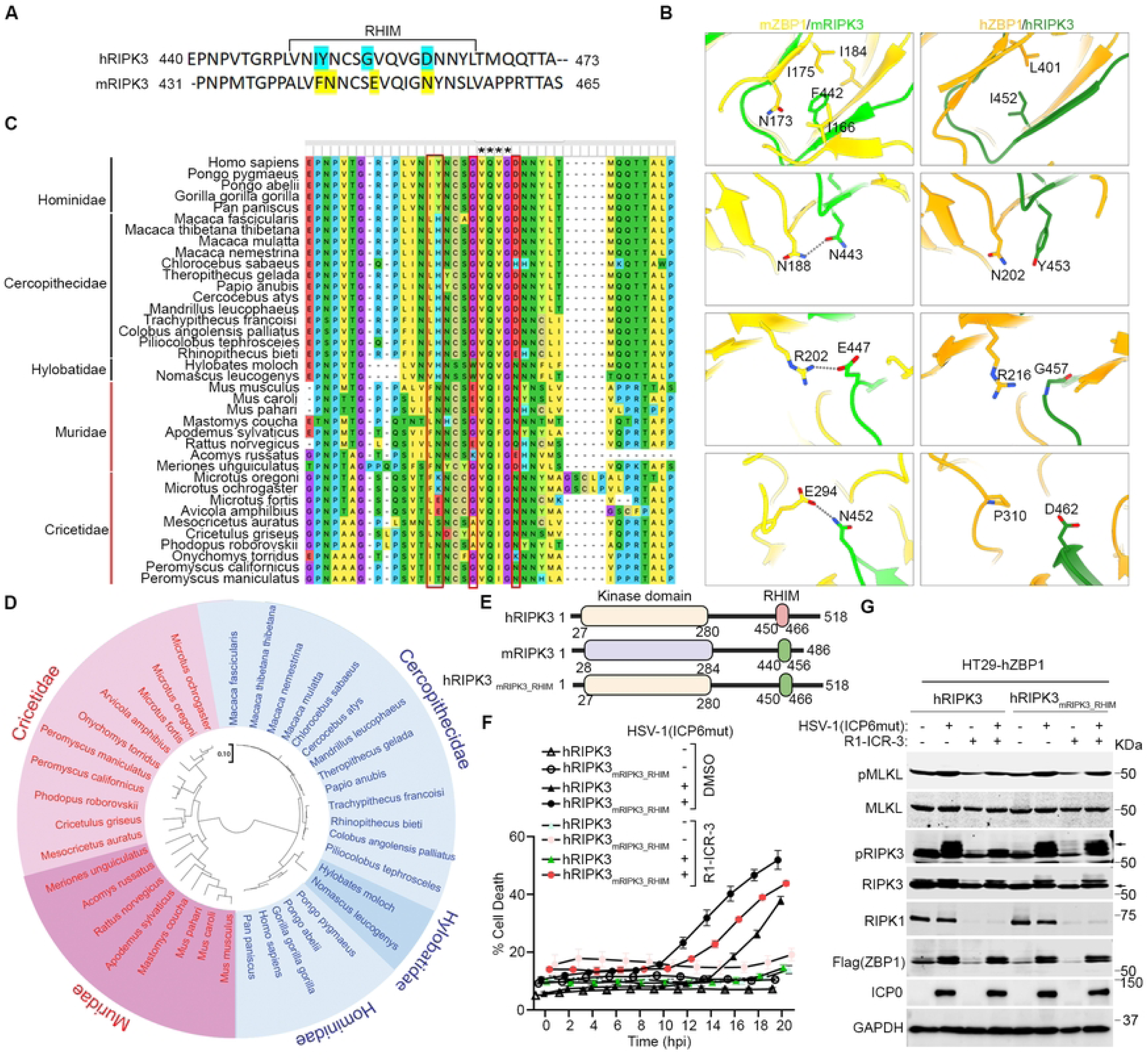
Species-Specific RIPK3 RHIM domain determines the requirement of RIPK1 in HSV-1(ICP6mut)-triggered ZBP1-dependent necroptosis. (A) Alignment of RHIM sequences from human RIPK3 (hRIPK3) and mouse RIPK3 (mRIPK3). Several variable residues between hRIPK3 and mRIPK3 analyzed below were highlighted. The alignment was performed using ClustalW2 online software. (B) Structural comparison between mZBP1-mRIPK3 and hZBP1-hRIPK3 complexes. mZBP1 was colored in yellow, mRIPK3 in light green, hZBP1 in orange, and hRIPK3 in dark green. Hydrogen bonds were indicated by gray dotted lines. (C) Multiple sequence alignment of RIPK3 protein sequences from Primate and Rodent species. The conservative RHIM core amino acids were highlighted with a star (*). The variable residues in (A) were highlighted with a red box. (D) Phylogenetic tree of RIPK3 proteins by MEGA11. RIPK3 protein sequences from Primate and Rodent species were retrieved from GeneBank. The evolutionary history was inferred using the Maximum Likelihood method. All sequences from Primates were highlighted with Blue, and all sequences from Rodents were highlighted with Red. (E) Schematic of hRIPK3, mRIPK3, and chimeric hRIPK3 construct-hRIPK3_mRIPK3_RHIM_. (F) Cell death kinetics of HT29-hZBP1 (RIPK3 KO) cells reconstituted with hRIPK3, and hRIPK3_mRIPK3_RHIM_ pretreated with DMSO or R1-ICR-3 (100 nM) for 5 hrs, followed by HSV-1(ICP6mut) infection. (G) Immunoblotting of HT29-hZBP1 (RIPK3 KO) cells reconstituted with hRIPK3, and hRIPK3_mRIPK3_RHIM_ pretreated with DMSO or R1-ICR-3 (100 nM) for 5 hrs, followed by HSV-1(ICP6mut) infection and subjected to immunoblotting with p-MLKL, MLKL, p-RIPK3, RIPK3, RIPK1, Flag (ZBP1), ICP0, and GAPDH antibodies. Results are representative of at least two independent experiments. Error bars represent mean ± SD.

The four residue variations (I452, Y453, G457, and D462) mentioned above are fully conserved within *Hominidae*, and G457 and D462 are conserved across all primates (Fig. 6C). Conversely, the four residue variations (F442, N443, E447, and N452) identified in the house mouse RIPK3 are highly conserved among the *Mus* and *Rattus* genera (Fig. 6C). Additionally, N452 is highly conserved among all rodent species (Fig. 6C). To understand the evolutionary relationships of RIPK3 among primate and rodent species, we performed a phylogenetic analysis using the maximum likelihood method in MEGA11. The analysis divided RIPK3 into two groups between primate and rodent species, with rodent species showing greater diversity compared to primate species (Fig.6D). Importantly, all the conserved RHIM sequences identified through sequence alignment clustered together (Fig.6D). Taken together, these sequence differences between humans and mice, and between rodents and primates in general, likely contribute to stronger binding between mZBP1 and mRIPK3 compared to their human counterparts, thus obviating the need for mRIPK1 recruitment in the mZBP1-mRIPK3 complex.

To investigate if these amino acid differences between the RHIMs of human and mouse RIPK3 dictated the species-specific requirement of RIPK1 in ZBP1 signaling in humans, we generated a chimeric construct, hRIPK3_mRIPK3_RHIM_, by swapping the RHIM of hRIPK3 with that of mRIPK3 (Fig. 6E). We then ablated endogenous RIPK3 in HT-29 cells (HT-29 RIPK3 KO) and reconstituted these cells with either wild-type hRIPK3 or an hRIPK3 with its RHIM swapped out for the murine equivalent. After establishing that HT29 (RIPK3 KO) cells expressing hRIPK3_mRIPK3_RHIM_ exhibited a comparable ability to mediate HSV-1(ICP6mut)-triggered necroptosis compared to cells expressing wild-type hRIPK3, we examined if hRIPK1 was now needed for hRIPK3_mRIPK3_RHIM_-driven signaling downstream of ZBP1. Remarkably, ablating RIPK1 (using the degrader) R1-ICR-3 did not affect HSV-1(ICP6mut)-triggered necroptosis in HT29-hRIPK3_mRIPK3_RHIM_ cells, demonstrating that RIPK1 is no longer necessary for hRIPK3 activation by ZBP1 when hRIPK3 harbors the RHIM from mRIPK3 (Fig. 6F, G).

### RIPK1 facilitates ZBP1-RIPK3 to form a functional amyloid signaling complex

Recent structural analysis has revealed that the RHIM domains of RIPK1 and RIPK3 mediate the formation of amyloid structures through hetero- or homo-oligomerization [37-39]. Purified ZBP1-RIPK1 and ZBP1-RIPK3 proteins have been demonstrated to undergo amyloid formation [37, 40], mirroring the behavior observed in RIPK1-RIPK3 complex. Here, we aimed to investigate the presence of amyloid formation during HSV-1(ICP6mut)-triggered ZBP1-driven necroptosis to shed light on the unique requirement for ZBP1-RIPK3 complex formation in human cells. Amyloids formed by RHIMs were classically characterized and quantified using aromatic, cross β binding dyes-Thioflavin T (ThT) [37, 39, 40]. We isolated the ZBP1-RIPK3 complex from HSV-1(ICP6mut)-infected HT29-hZBP1 cells, or from HT29-hZBP1 (RIPK1 KO) cells for subsequent testing of ThT binding. While the complex from HT29-hZBP1 cells displayed a distinct ThT fluorescence pattern from controls, the ZBP1-containing complex obtained from RIPK1 KO cells exhibited a similar ThT fluorescence pattern as the buffer alone- or ThT dye alone control groups (Fig. 7A). This finding supports the idea that RIPK1 is essential for the formation of functional ZBP1-RIPK3 amyloid complexes.

**Figure 7.**
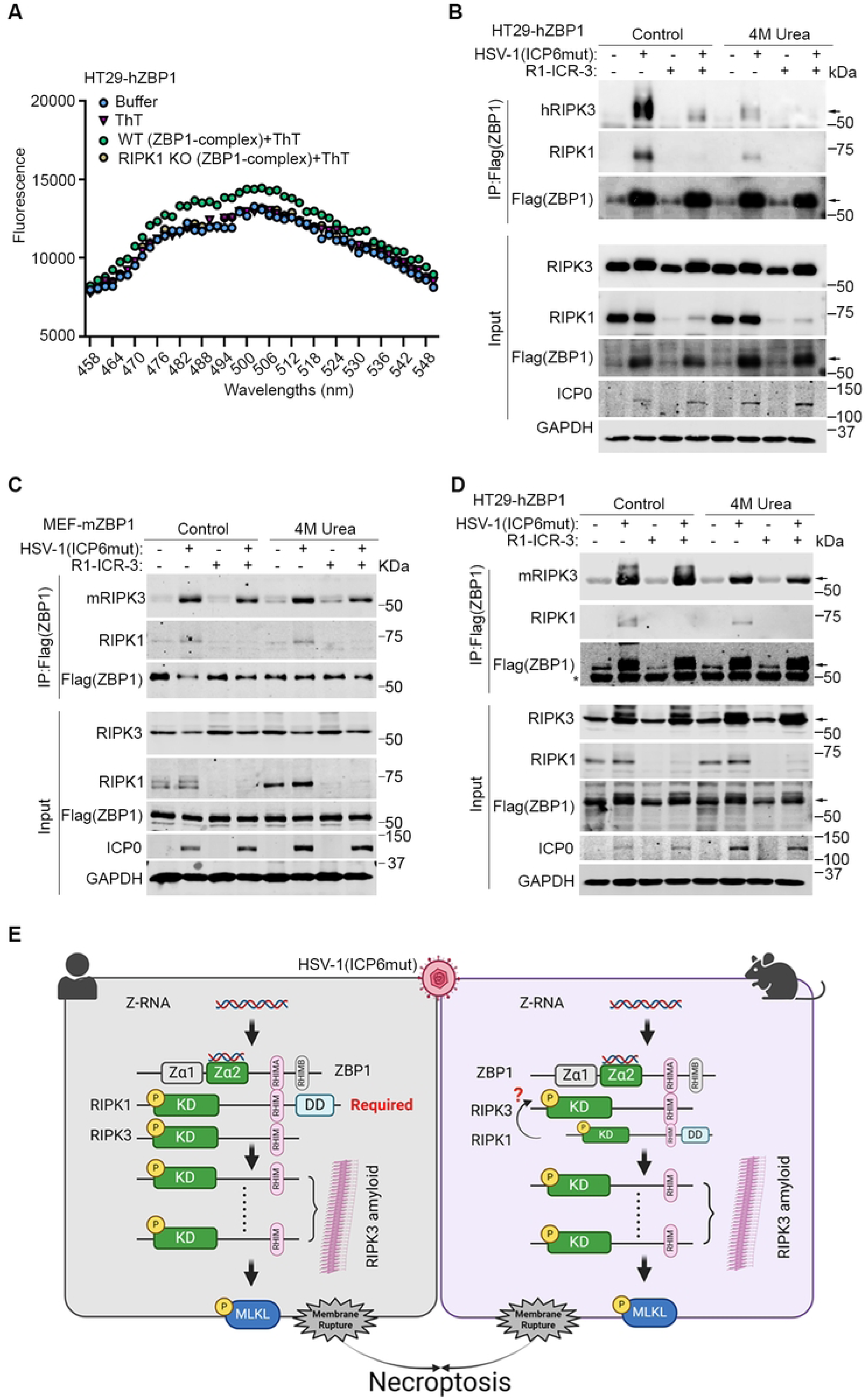
RIPK1 facilitates the formation of a ZBP1-RIPK3 amyloid structure in human cells. (A) Thioflavin T (ThT) binding assay of endogenous ZBP1-containing complexes from HSV-1(ICP6mut) infected HT29-hZBP1 and HT29-hZBP1 (RIPK1 KO) cells. (B) HT29-hZBP1 (RIPK3 KO) cells transduced with human RIPK3 (hRIPK3) were either mock or HSV-1(ICP6mut) infection for 10 hrs, with or without R1-ICR-3 (100 nM). Cells were harvested and subjected to immunoprecipitation. ZBP1 complexes isolated by immunoprecipitation were lysed in regular lysis buffer or buffer containing 4M urea. (C) MEF-mZBP1 cells were either mock or HSV-1(ICP6mut) infection for 10 hrs, with or without R1-ICR-3 (1 µM). Cells were harvested and subjected to immunoprecipitation. ZBP1 complexes isolated by immunoprecipitation were lysed in regular lysis buffer or buffer containing 4M urea. (D) HT29-hZBP1 (RIPK3 KO) cells reconstituted with mouse RIPK3 (mRIPK3) were either mock or HSV-1(ICP6mut) infection for 10 hrs, with or without R1-ICR-3 (100 nM). Cells were harvested and subjected to immunoprecipitation. ZBP1 complexes isolated by immunoprecipitation were lysed in regular lysis buffer or buffer containing 4M urea. (E) Schematic model of the proposed ZBP1-driven necroptosis in both human and mouse cells during HSV-1 (ICP6mut) infection. Results are representative of at least two independent experiments. Error bars represent mean ± SD.

As amyloid structures are resistant to disassembly by urea [37], we treated the ZBP1-complex isolated from HT29-hZBP1 cells treated with or without R1-ICR-3, followed by 4M urea treatment. The ZBP1-containing complex isolated from HT29-hZBP1 (RIPK3 KO) cells reconstituted with hRIPK3 still preserved when treated with 4M urea, while failed in the R1-ICR-3-treated group (Fig.7B). We then explored this mechanism in mouse cells. The ZBP1-complex of MEF-mZBP1 cells was immunoprecipitated and subjected to 4M Urea treatment. As shown in Fig. 7C, mZBP1-mRIPK3 complex formation is independent of RIPK1 and more resistant to 4M Urea treatment compared to hZBP1-hRIPK3 complex.

To confirm whether the dependency on RIPK1 is due to species-specific differences in RIPK3, we reconstituted HT29-hZBP1 (RIPK3 KO) cells with mouse RIPK3. Similar to MEF-mZBP1 cells, hZBP1-mRIPK3 complex formation during viral infection is independent of RIPK1 and resistant to 4M treatment (Fig.7D), indicating that mRIPK3 tends to form a more stable complex with ZBP1 compared to hRIPK3. These results indicate that during HSV-1(ICP6mut) infection, mRIPK3 efficiently binds to activated ZBP1, forming a stable amyloid structure. Conversely, hRIPK3 relies on RIPK1 to facilitate the assembly of a stable ZBP1-RIPK3 amyloid complex.

## Discussion

During virus-triggered ZBP1 activation, it is well-known that the heterotypic RHIM-RHIM interactions between ZBP1 and RIPK3 initiates RIPK3 activation and cell death [29]. The mechanism is not limited to large DNA viruses including MCMV, HSV, and poxvirus VACV, but also extends to the RNA viruses IAV, and SARS-CoV-2 [24, 25]. Here, we show that RIPK1 is crucial for ZBP1-driven cell death triggered in human cells during HSV-1(ICP6mut) infection, where it facilitates the assembly of a stable ZBP1-RIPK3 amyloid complex (Fig.7E). In mouse cells, ZBP1 can bind RIPK3 and activate downstream necroptosis signaling without the need for RIPK1 (Fig.7E).

Replacement of human RIPK3 with mouse RIPK3 (or even just its RHIM) in human cells allows the formation of a stable ZBP1-RIPK3 complex without the need for RIPK1, suggesting that mouse RIPK3 exhibits a higher binding affinity for ZBP1, compared to human RIPK3. Indeed, mouse RIPK3 can compete with human RIPK3 for binding to ZBP1, but not vice versa (data not shown). When necroptosis is triggered in human cells, the low binding affinity of RIPK3 necessitates the presence of RIPK1 to facilitate ZBP1-RIPK3-mediated necroptosis. In other words, human ZBP1 binds to RIPK1 independent of RIPK3, and the ZBP1-RIPK1 interaction is necessary to form and stabilize a functional ZBP1-RIPK3 signalosome. Chemical inhibition of RIPK1 kinase activity, or transient degradation by R1-ICR-3 each prevents HSV-1(ICP6mut)-triggered ZBP1-driven necroptosis only in human cells. In contrast, in mouse cells, chemical inhibition of RIPK1 kinase activity only delayed virus-induced cell death, while R1-ICR-3-mediated degradation had little impact on ZBP1-initiated necroptosis. These results suggest that in mouse cells, the kinase activity of RIPK1 may contribute to RIPK3 activation, while the recruitment of RIPK3 is not dependent on RIPK1 for the formation of ZBP1-RIPK3 oligomers, consistent with previous reports system [41].

We identified four residue variants that may contribute to the differences in ZBP1-RIPK3 binding affinity between the two species. These residues (I452, Y453, G457, and D462) in human RIPK3 are conserved between Hominids. Notably, G457 and D462 are conserved across all primate species. Among the Cercopithecidae and Hylobatidae families, I452 and Y453 have been replaced with L452 and H453, respectively. Based on structural analysis, L452 is likely to exhibit similarities to I452 in human RIPK3. It may form hydrophobic interactions with an additional amino acid residue in hZBP1, although not to the same extent as mediated by F442 of mRIPK3 due to its smaller side chain. Similarly, H453 is unlikely to form a hydrogen bond with N202 in hZBP1 due to its shorter side chain and the resulting longer distance to the side chain of N202. N443 and N452 of mouse RIPK3 are highly conserved among rodent species. Although some species have Glu (E), Lys (K), and Thr (T) at position 443 in RIPK3, all these three residues have the potential to engage in hydrogen bond formation with N188 of mZBP1, as shown in Figure 6C.

We further extended our observations on the species-specific requirement for RIPK1 in ZBP1-mediated activation of RIPK3 to include orthomyxoviruses (IAV), poxviruses (VACV), and the ZBP1-activating compound CBL0137 [42] (data not shown). These findings underscore the essential and universal role of RIPK1 in promoting the formation of the ZBP1-RIPK3 complex and consequent cell death in human cells. This highlights the potential of RIPK1 degraders or kinase inhibitors as promising therapeutics in ZBP1 signaling-mediated pathogenesis in human diseases. ZBP1-initiated necroptosis contributes to severe inflammatory tissue damage, such as those caused by the influenza virus and SASR-CoV-2 [20, 21, 24, 30]. Inhibitors of RIPK3 kinase activity prevent lung injury in mouse models of severe influenza [30]. Our unanticipated finding that ZBP1-mediated necroptosis in humans (but not mice) requires RIPK1 now positions existing RIPK1 kinase inhibitors as effective therapeutics for ZBP1-initiated pathologies, including but not limited to, viral diseases such as influenza-driven Acute Respiratory Distress Syndrome. Notably, several RIPK1 kinase inhibitors have already entered clinical trials [43], making this strategy feasible.

It remains to be defined whether ZBP1-RIPK1 signaling contributes to TLR3/4-induced inflammatory signaling in human cells, as it does in mouse cells [44]. It is also unclear if RIPK1 is required for ZBP1-RIPK3 complex formation and/or ZBP1-driven immunopathology in human Aicardi-Goutiere’s patients, where ZBP1 activation contributes to cell death, fatal autoinflammation, and immune pathology [42, 45-47].

In summary, our findings demonstrate an essential and universal role for RIPK1 in promoting the formation of the ZBP1-RIPK3 complex and consequent cell death in human cells, highlighting the potential of RIPK1 degraders or kinase inhibitors as potential therapeutics in ZBP1 signaling-mediated pathogenesis in human diseases.

## ACKNOWLEDGMENTS

We thank Dr. Jason Bodily for the supply of primary human foreskin fibroblasts (HFF). We thank Dr. Edward Mocarski and Ann E. McDermott for critical reading and discussion of the manuscript. We acknowledge the INLET Core facility of LSU Health Shreveport for their assistance with Incucyte Zoom live cell imaging, and the CAIPP Bioinformatics and Modeling Core for modeling analyses. This work is supported by the NIH R21 R21AI175590 and NIH C0BRE P20GM134974 Pilot Grant to H.G. Work in the P.M. lab is funded by Breast Cancer Now as part of Programme Funding to the Breast Cancer Now Toby Robins Research Centre (CTR-QR14-007) and CRUK Programme Funding (C26866/A24399). The Meier lab acknowledges NHS funding to the NIHR Biomedical Research Centre. Work in the S.B. lab is supported by NIH grants AI135025, AI168087, AI144400, and AI161624, and by NIH Cancer Center Support Grant P30CA006927. Work in the H.K. lab is supported by PG00020481.

## AUTHOR CONTRIBUTIONS

O.A., S.W., and H.G. carried out most of the experiments, with assistance from Y.C., S.P., J.Y. C.Y., and T.Z. H.K. performed experiments and edited the manuscript. C.L. and J.W. carried out AlphaFold analyses. T.T., and R.W. helped with RIPK1 PROTAC and RIPK1 reconstitution experiments. B.B. developed the RIPK1 PROTAC. J.U. provided resources and edited the manuscript. P.M. provided resources and guidance in support of experiments. S.B. provided resources and guidance in support of experiments, and edited the manuscript. H.G. was responsible for the overall design of the study, performed experiments, prepared figures, and wrote the manuscript.

## Materials and Methods

### Cells and Viruses

Human colorectal adenocarcinoma cell line (HT29) expressing Flag-hZBP1 as well as mutants, Immortalized MEFs expressing Flag-mZBP1 as well as mutants, immortalized human foreskin fibroblasts (HS68), and primary human foreskin fibroblasts (HFF), HEK 293T cells were maintained in DMEM containing 4.5 g/ml glucose, 10% fetal bovine serum (Sigma), 2 mM L-glutamine, 100 units/ml penicillin, and 100 units/ml streptomycin (Invitrogen). HT29, responding efficiently to express endogenous ZBP1 upon human IFN-β priming, was kindly provided by Dr. Heather Koehler (Washington State University) and cultured with McCoy’s 5A medium supplemented as above. U937 cells were grown in RPMI1640 with the above supplements. HFF cells were isolated from discarded foreskin circumcisions as previously described [48] and kindly provided by Dr. Jason Bodily (LSUHSC-Shreveport).

The ICP6 RHIM mutant of HSV1 (HSV-1(ICP6mut)) was used as reported before [49] at an MOI of 5 unless otherwise specified. Viral stocks were propagated and tittered on monolayer cultures of Vero cells. IAV [27] and VACV [28] propagation and infection were performed as previously described.

### Generation of knockout and reconstitution of cell lines

MEF cells of ZBP1 knockout were generated using CRISPR/Cas9-mediated ablation as described before [20], and ZBP1 knockout cells were reconstituted with Flag-mZBP1 as well as mutants via pQCXIH retroviral vector (Clontech) and retrovirus as previously described [19]. HT-29 cells of RIPK3 knockout (KO), RIPK1 KO, or ZBP1 KO were generated by using CRISPR/Cas9 gene editing technique [50]. In brief, the gRNAs targeting the human RIPK1, RIPK3, or ZBP1 locus were designed using the web tool CRISPOR (http://crispor.org). The gRNAs with the highest score and the least potential off-target activities were selected. The Cas9-target sites are listed as follows: human RIPK3: 5’-CTCGTCGGCAAAGGCGGGTT-3’; human RIPK1: 5’-CTCGGGCGCCATGTAGTAG-3’; human ZBP1: 5’-CGGTAAATCGTCCATGCTT-3’ (Sigma). The gRNAs were cloned into pSpCas9n(BB)-2A-GFP plasmid (PX461, Addgene). HT29 cells were transfected with these gRNAs and sorted onto 96-well plates via flow cytometry to form single cell clones. The KO cells were confirmed by the immunoblotting assay. RIPK3 KO cell lines were reconstituted with Wild-type (WT) human RIPK3, and mutants as previously described [36]. HT29-hZBP1 (RIPK1 KO) cells, reconstituted with either human RIPK1 or mouse RIPK1, as well as non-reconstituted cells, were kindly provided by Dr. Pascal Meier. In brief, HT29 parental cells were transfected with pMA-T-gRNA-RIP1-3 (synthetic construct expressing the guide) targeting RIPK1 and Cas9-GFP (pSpCas9(BB)-2A-GFP (PX458, Addgene). The Cas9-target sites are: 5’-GCTCGGGCGCCATGTAGTAG-3’. Cells were FACS (BD FACSymphony S6) sorted into single cell clones and analyzed for RIPK1 expression by immunoblotting using two separate (N-terminal and C-terminal) anti-RIPK1 antibodies. Cell death assays verified that the clones lacked functional RIPK1. RIPK1 was subsequently reintroduced with the Piggybac transposon system (PB-Cuo-MCS-IRES-GFP-EF1a-CymR-Puro Cumate inducible vector, PBQM812A-1, System Biosciences), and cells were selected with puromycin 1µg/ml for 7-10 days. RIPK1 expression was induced with water-soluble Cumate solution (QM150A, System Biosciences) for the times indicated.

### siRNA electroporation transfection of primary fibroblast cells

Electroporation transfection of *siZbp1* was performed on HFFs in a Lonza 4D-NucleofectorTM X unit (Lonza, Cat#: AAF-1003X). Briefly, HFFs were maintained in an antibiotics-free DMEM medium with 10% FBS for 24 hours before electroporation. Cells then were washed with PBS, harvested with Trypsin-EDTA, suspended in PBS, and counted. 1x10^6^ HFFs were transfected with 50 nM siZBP1 Smartpool (L-014650-00-0010, Dharmacon, ON-TARGETplus Human ZBP1 siRNA-SMARTpool) or the same amount of negative control siRNA (D-001810-10-05, Dharmacon, ON-TARGETplus Non-targeting Control pool) in 100 µl transfection buffer, following the protocols provided by the manufacturers using the Basic Nucleofector™ Kit for Primary Mammalian Fibroblasts (VPI-1002, Lonza). After transfection, HFFs were cultured with or without IFN-β for 24 hours. 5x10^5^ HFFs were collected for immunoblotting assay and the rest of the cells were subjected to mock- or virus-infection in 96-well plates for cell viability assay. The siZBP1 SMARTpool target sequence are: seq1: 5’-CAAAGUCAGCCUCAAUUAU-3’; seq2: 5’-GGAUUUCCAUUGCAAACUC-3’; seq3: 5’-GGACACGGGAACAUCAUUA -3’; seq4: 5’-CAAAAGAUGUGAACCGAGA -3’.

### Cell Viability Assay

Viability was assessed with a SYTOX nuclear stain exclusion assay or by a CellTiter-Glo Luminescent Cell Viability Assay (G7573, Promega). Relative viability is calculated by dividing the three technical replicates for each treatment condition by the average of measurement luminescence values of the reference condition as specified for each figure (such as uninfected group or untreated group). The individual data points indicate three technical repeats for each condition. Results are representative of at least 2 independent experiments. For SYTOX nuclear stain exclusion assay, cell death kinetics were quantified in real-time by using an Incucyte Zoom Live Cell Imaging and Analysis (Sartorius). Briefly, plated cells were incubated in media containing the specified treatments plus 50 nM of the membrane impermeant dye SYTOX Green (S7020, Invitrogen). Two images were collected per well of 96-well plates (3603, Corning) unless otherwise specified. SYTOX^+^ cells were considered dead and quantified using Incucyte image analysis software. The total cell count for each image was determined at the endpoint. The percentage (%) value of cell death, representing cell death kinetics, was calculated by normalizing the dead cell count to the total cell count respectively.

### Cell Treatments

Inhibitors including GSK963 (S8642, Selleckchem, 5μM), GSK872 (S8465, Selleckchem, 5μM), Nec-1 (S8037, Selleckchem, 30μM), Necrosulfonamide (NSA, 100573, MedChemExpress, 1μM) or zVAD-fmk (S7023, Selleckchem, 25μM) were applied to cells 30 mins prior to and during viral infection. R1-ICR-3, kindly provided by Dr. Pascal Meier, was applied to cells 5 hrs prior to and during viral infection. siRNA transfection: cells were plated in 96-well or 60mm-dish at 60% confluency, allowed to adhere for 24 hrs and transfected with Silencer Select Pre-Designed siRNAs (s72975 for murine si*Ripk1*, Thermo Fisher) or control siRNA (4390843, Thermo Fisher) using RNAiMAX transfection reagent (13778075, Thermo Fisher) according to manufacturer’s recommendations. Following transfection, cells were incubated for 48 hrs, either subjected to mock- or virus-infection for 10 hrs for Immunoblotting or 18 hrs for cell viability assay.

### Plasmid/Retroviral Vector Constructs and Transduction

To create hZBP1-expressing retroviral vector, three-tandem FLAG epitope-tagged hZBP1 as well as mutants open reading frame (ORF) were inserted into pQCXIH retroviral vector (Takara Bio). Overlap extension PCR was employed to generate expression constructs of hZBP1 mutants, tetra-Ala RHIM A domain (amino acids 206-209) substitution (mutRHIMA), tetra-Ala RHIM B domain (amino acids 264-267) substitution (mutRHIMB), N46D/Y50A (mutZα), N141D/Y145A (mutZβ), and N46D/Y50A plus N141D/Y145A (mutZαmutZβ). For RHIM swapping experiments, MYC epitope-tagged hRIPK3 or hRIPK3_mRIPK3_RHIM_ ORF were generated by Twist Bioscience and inserted into pQCXIH retroviral vector. All plasmids were verified by DNA sequencing. Retrovirus stock was prepared from 293T cells that were transfected with retrovirus constructs along with pCMV-Tat, psPAX2, and VSV-G-expressing plasmids [19]. HT29 were transduced with retrovirus and selected with 500μg/ml hygromycin (Thermo Fisher), respectively.

### Immunoprecipitation and Immunoblotting

Cell pellet was collected and re-suspended in lysis buffer (25 mM Tris-HCl pH 7.4, 150 mM NaCl, 1 mM EDTA, 1% NP-40 and 5% glycerol, Roche complete protease inhibitor, and phosphatase inhibitor). Cell lysates were incubated on ice for 10 min, and briefly sonicated to shear chromatin, then cleared by high-speed centrifugation (13,000rpm, 15min) at 4°C. After saving 5% of the total cell lysate for input, the extracts were subjected to immunoprecipitation with anti-FLAG M2 magnetic beads (Sigma, M8823) overnight at 4 °C. The next day, beads were washed with lysis buffer and the immunoprecipitants were eluted off the beads using SDS-PAGE sample buffer. The supernatants were subjected to immunoblot analysis. Primary antibodies were used at the following dilutions: phosphorylated human MLKL (91689, Cell Signaling, 1:1000), phosphorylated murine MLKL (37333, Cell Signaling, 1:1000), total MLKL (MABC604, Sigma, 1:1000), RIPK1 (4926, Cell Signaling, 1:1000), RIPK1 (610459, BD Transduction Laboratories™, 1:1000), phosphorylated RIPK1 (44590, Cell Signaling, 1:1000), human RIPK3 (13526, Cell Signaling, 1:1000), phosphorylated RIPK3 (93654, Cell Signaling, 1:1000), murine RIPK3 (2283, ProSci, 1:1000), FLAG (20543, Proteintech, 1:2000), ICP0 (sc-53070, Santa Cruz, 1:500), GAPDH (60004, Proteintech, 1:4000).

### Thioflavin T (ThT) Binding Assay

2×10^8^ HT29-hZBP1 or HT29-hZBP1(RIPK1KO) cells were infected with HSV-1(ICP6mut) (MOI=2) for 10 hrs. ZBP1 complex was purified by anti-FLAG M2 magnetic beads. The resulting protein complex was eluted from the beads with low pH elution buffer (21004, Thermo Scientific), followed by neutralization with 1/10 volume of 1 M Tris-HCl (pH 8.0). For ThT fluorimetry, 50 μl ZBP1 complexes were incubated with 50 μM ThT (ab120751, Abcam) for 30 min at room temperature. ThT fluorescence was measured using The Spark Multimode Microplate Reader (Tecan). Excitation was performed at 430 nm and emission was measured from 450 to 550 nm.

### Prediction of ZBP1-RIPK3 Interaction Model by AlphaFold

The AlphaFold structures in this study were mainly generated from the AlphaFold2 [51, 52] implementation in DGX-A100 HPC(8 Ampere NVIDIA A100 40Gb 1215MHz GPUs, 6 AMD EPYC 7742 64 Core 2.25GHz Processors) from the department of Psychiatry and Behavioral Medicine, LSUHSC-shreveport, using the default settings with Amber relaxation (msa_method=mmseqs2, homooligomer=1, pair_mode=unpaired, max_msa=512:1024, subsample_msa=True, num_relax=5, use_turbo=True, use_ptm=True, rank_by=pLDDT, num_models=5, num_samples=1, num_ensemble=1, max_recycles=3, tol=0, is_training=False, use_templates=False). AlphaFold was run once with each of the 5 trained models; the five models generated were checked for consistency, and unless specified otherwise, the top-ranked model was taken in each case for density fitting. AlphaFold computes pLDDT score and pTM score to indicate the accuracy of a prediction. For complex prediction, sequences were entered in tandem and separated by a semicolon. The top-ranked model was taken in each case for density fitting. We used pTM for ranking protein-protein complexes. The structural models generated by AlphaFold were aligned and analyzed in Coot [53]. The molecular representations were prepared using UCSF ChimeraX [54].

### Statistical Analysis

Statistical significance was determined by use of either Student’s t-test or One-way ANOVA. Statistical analysis was performed in GraphPad Prism9 (version 9.1.1). Statistical significance is indicated as follows: *P < 0.05, **P < 0.01, ***P < 0.001, ****P < 0.0001 in the corresponding figures.

## SUPPLEMENTAL LEGENDS

**Figure S1. Related to Figures 2 and 3.** (A-F) Cell death kinetics of SVEC4-10 (A), 3T3-SA (C), and MEF-mZBP1 (E) cells treated with DMSO or R1-ICR-3 (1µM) for 5 hrs, followed by HSV-1(ICP6mut) infection. Immunoblotting of SVEC4-10 (B), 3T3-SA (D), and MEF-mZBP1 (F) cells pretreated with DMSO or R1-ICR-3 (1µM) for 5 hrs, followed by HSV-1(ICP6mut) infection and subjected to immunoblotting with p-MLKL, MLKL, RIPK1, ICP0, and GAPDH antibodies. (G) Cell death kinetics of MEF-mZBP1, followed by HSV-1(ICP6mut) infection, with or without Nec-1, GSK963, GSK872, NSA, or zVAD. (H) MEF reconstituted with empty vector (EV) or Flag-mZBP1, were either mocked or infected with HSV-1(ICP6mut) for 10 hrs. Co-immunoprecipitation was performed in these cells. Results are representative of at least two independent experiments. Error bars represent mean ± SD.

**Figure S2. Related to Figure 5.** (A) Cell viability of MEFs reconstituted with the indicated mouse ZBP1 constructs, followed by HSV-1(ICP6mut) infection. Viability was determined at 18 hpi by CellTiter-Glo assay. (B) Cell death kinetics of MEFs reconstituted with mZBP1 mutants, followed by HSV-1(ICP6mut) infection. (C) The expression levels of mZBP1 in MEFs reconstituted with mZBP1 constructs were confirmed by immunoblotting with Flag and GAPDH antibodies. One-way ANOVA and Dunnett’s multiple comparisons tests were used to test for statistical differences in (A). *P < 0.1, **P < 0.0001, ***P < 0.001, ****P < 0.0001. Individual data points indicate three technical replicates. Results are representative of at least two independent experiments. Error bars represent mean ± SD.

